# Predictive Coding Explains Asymmetric Connectivity in the Brain: A Neural Network Study

**DOI:** 10.1101/2025.02.27.640572

**Authors:** Romesa Khan, Hongsheng Zhong, Shuvam Das, Jack Cai, Matthias Niemeier

## Abstract

Seminal frameworks of predictive coding propose a hierarchy of generative modules, each attempting to infer the neural representation of the module one level below; the predictions are carried by top-down feedback projections, while the predictive error is propagated by reciprocal forward pathways. Such symmetric feedback connections support visual processing of noisy stimuli in computational models. However, neurophysiological studies have yielded evidence of asymmetric cortical feedback connections. We investigated the contribution of neural feedback during sensorimotor processes, in particular visual processing during grasp planning, by utilizing convolutional neural network models that had been augmented with predictive feedback and were trained to compute grasp positions for real-world objects. After establishing an ameliorative effect of symmetric feedback on grasp detection performance when evaluated on noisy stimuli, we characterized the performance effects of asymmetric feedback, similar to that observed in the cortex. Specifically, we tested model variants extended with *short*-, *medium*- and *long*-range feedback connections (i) originating at the same source layer or (ii) terminating at the same target layer. We found that the performance-enhancing effect of predictive coding under adverse conditions was optimal for *medium*-range asymmetric feedback. Moreover, this effect was most prominent when *medium*-range feedback originated at a level of representational abstraction that was proximal to the input layer, in contrast to more distal layers. To conclude, our simulations show that introducing biologically realistic asymmetric predictive feedback improves model robustness to noisy visual stimuli in a neural network model optimized for grasp detection.

**Significance statement:** It is commonly held that the brain predicts the causes of its sensorium via top-down neural pathways. While canonical models of predictive coding assume reciprocal feedforward and feedback connections, functional evidence highlights the importance of non-reciprocal ‘asymmetric’ feedback, whose role remains poorly understood, particularly in sensorimotor functions. Using neural network models of grasp planning, we characterized optimal pathlengths and source regions for asymmetric feedback facilitating visuomotor processing of noisy sensory inputs. Our findings show that *medium*-range feedback from early layers marks a sweet spot, incorporating optimal distance between the neural representations of source/target layers and representational abstraction of the feedback source. This intimates an uncharted role of intermediate brain areas along the visuomotor stream as a source of predictive signals.

## Introduction

Descending neural pathways transmit neural signals from higher cortical areas back to earlier processing stages, enabling top-down modulation of lower-level neural circuits. Such feedback has been recognized as central for perceptual and cognitive processes, enabling the brain to implement learning mechanisms (Roelfsema & Holtmaat, 2018), attentional modulation (Baluch & Itti, 2011; Desimone & Duncan, 1995) and refinement of sensory representations based on context, expectations, and prior experience (Gilbert & Li, 2013; Kok, Jehee, et al., 2012; Mumford, 1992). Crucially, feedback is believed to be instrumental for conveying predictions.

That is, the brain has been widely conceptualized as a “prediction machine” that relies on internal generative models to actively construct explanations for the causes of noisy sensory inputs (Friston, 2005). Specifically, it is held that predictive feedback signals are conveyed via cortico-cortical top-down projections that are then compared to bottom-up signals such that the deviations from these signals, in the form of predictions errors, are carried forward. An influential framework of such predictive coding proposes a hierarchy of generative modules, each attempting to infer the neural representation of the module below, with topographically reciprocal feedforward connections (Lee & Mumford, 2003; Pennartz et al., 2019; Rao & Ballard, 1999a; Shipp, 2016). This idea maps well on previous anatomical data: Felleman and Van Essen (1991) proposed a hierarchical model of the visual cortex where adjacent areas often maintain both feedforward and feedback pathways, implying a measure of reciprocity in the network (Markov et al., 2011; Rockland & Pandya, 1979). Indeed, tracing studies in macaques reveal that reciprocal links between closely related visual areas significantly account for interareal communication (Markov et al., 2011). Such symmetric feedback connections are particularly important for object recognition under noisy or ambiguous viewing conditions. Hupé et al. (1998) have shown that area V2 modulates V1 responses, enhancing the detection of figures embedded in cluttered or noisy images. Furthermore, recurrent interactions between V1 and higher visual areas, including V2, are crucial for figure–ground segregation (Lamme & Roelfsema, 2000). Similarly, in computer simulations, symmetric feedback connections support object discrimination (Wen et al., 2018), particularly when processing noisy stimuli (Choksi et al., 2021; Huang et al., 2020).

However, there is growing evidence of asymmetric, non-reciprocated cortical feedback connections from quantitative mapping studies (Markov et al., 2014; Salin & Bullier, 1995). Long-range descending pathways cascade over multiple cortical areas (Kravitz et al., 2013; Rockland & Van Hoesen, 1994), and an advantage of long-latency over short-latency visuomotor feedback has been observed during cortical reward processing (Codol et al., 2023). Furthermore, experiments studying visuospatial attention in primates indicate a functional role of medium-range predictive feedback, extending from area V4 to area V1, in encoding accuracy of input stimuli (Debes & Dragoi, 2023), suggesting a potential benefit of *medium*-range feedback in signal processing under noise. This is similar to the previously suggested role of *medium*-range feedback in disambiguating signal from noise during global contour integration (Chen et al., 2014).

To investigate the contribution of asymmetrical neural feedback, here we simulated sensorimotor functions utilizing convolutional neural networks (CNNs) as a modelling framework. Hierarchical CNN architectures have been commonly used for object recognition tasks (Girshick et al., 2014) and are posited as suitable models of vision in the primate brain (Guclu & van Gerven, 2015; Yamins et al., 2014). Crucially, rather than solely relying on the feedforward flow of information in the canonical architecture of CNNs, we augmented them with symmetric, or asymmetric generative feedback loops that carried advanced representations to earlier layers of the networks, mimicking various aspects of feedback in the biological brain. The aim of the study was to understand (a) if an ameliorative effect of feedback can be isolated during action-guided visual tasks when the incoming signal is corrupted, and (b) to explore the layer-dependence of such predictive feedback in our model, thus shedding light on the relative functional contribution of cortico-cortical connectivity of asymmetric feedback originating at varying levels of representational abstraction.

## Methods

### Neural network architecture

We first trained two versions of a feedforward model, a short and a long backbone, on the task of grasp point generation. As a second step we then augmented the networks with feedback connections (feedback model) that conveyed predictive signals.

#### Short feedforward model

In detail, for step 1 we obtained the architecture of the short backbone model by modifying a custom convolutional neural network (CNN) architecture (see Figure 1(a)) into which we inputted photographic images with 4 channels (aside from red, green, and blue there was a depth channel, RGB-D, as an approximation of stereovision, that is known to be relevant for grasp performance (Watt & Bradshaw, 2000b)). The RGB-D images were passed into an input layer with two arms, specialized for receiving RGB and depth channels, respectively. The backbone contained a ‘feature compression’ module consisting of 4 convolution blocks that were connected through maximum pooling (MaxPool) layers to attain dimensionality reduction and invariant feature extraction (Simonyan & Zisserman, 2014). That is, each convolution block consisted of two or three convolution (Conv) layers, each of which was sandwiched together with a batch normalization (BN) and a rectified linear unit (ReLU) activation layer. Next, we added a ‘feature expansion’ module, composed of Conv and ConvTranspose2d layers (Paszke et al., 2019) that upscaled the feature maps back to the same spatial dimensions as the input (i.e., the entire network had an autoencoder style architecture). The output layer was a regression head that generated a 6-channel array of pixel-wise maps, each corresponding to one of six grasp parameters required for grasping (grasp centre coordinates (x, y); orientation; grasp opening; gripper size; grasp quality score), similar to (Kumra et al., 2022). The sole purpose of the feature expansion module together with its map-style output was not to attain biological realism but to facilitate interpolation of the infinite number of correct grasp solutions from the finite set of ground truth labels thereby facilitating training of the feedforward network (e.g., (Kumra et al., 2022)). The network architecture of the short backbone is detailed in Table 1.

**Figure 1.**
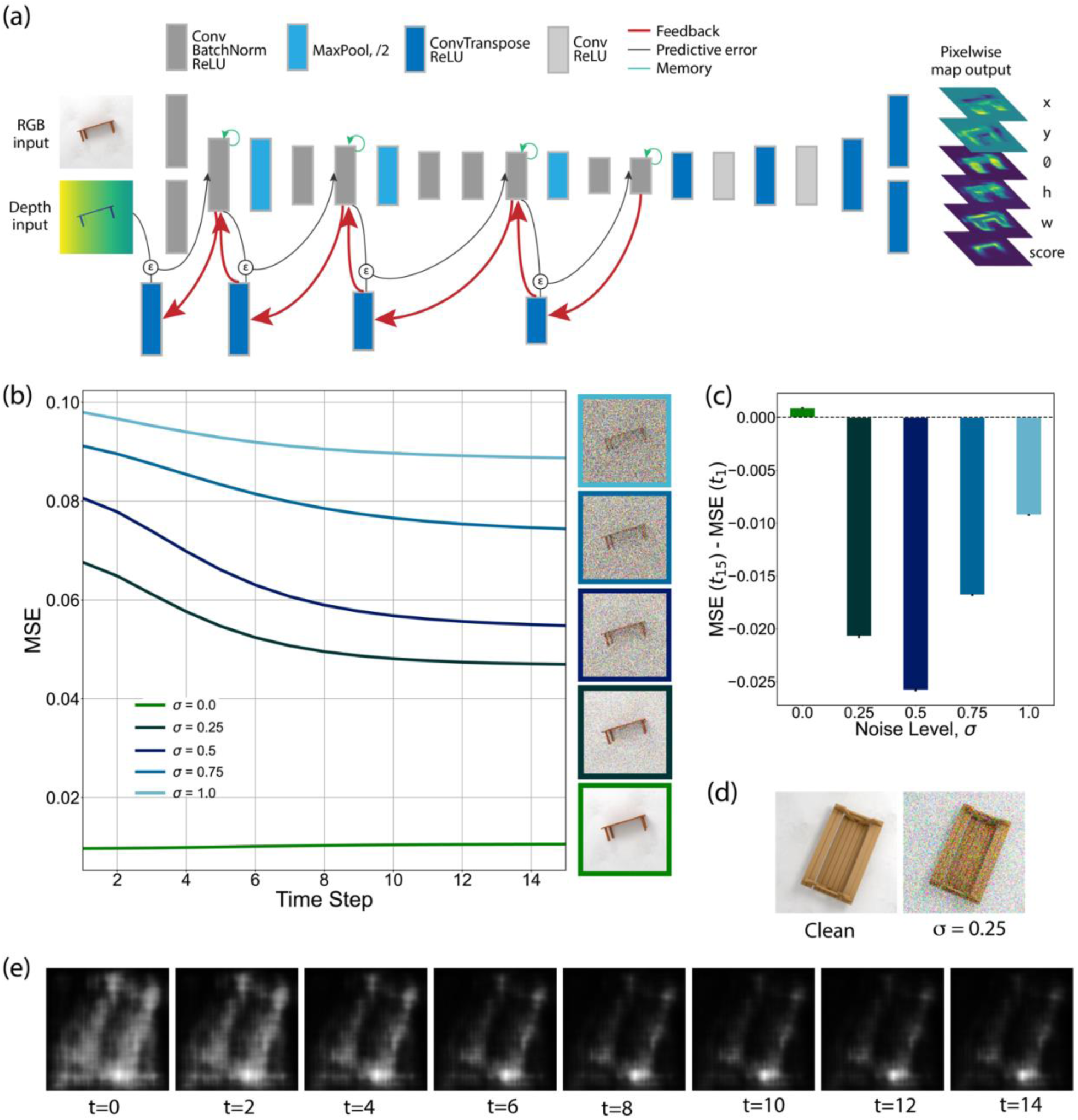
Predictive coding dynamics improve model performance for noisy stimuli. **(a)** Architecture of the grasping neural network, derived from the *short* feedforward backbone. Red arrows denote top-down feedback connections. Black arrows represent the error-correction process, whereby the predictive error (*ε*) drives the higher layer (feedback source) representation to better match the early layer representation (feedback target). Green arrows denote recurrent memory processes, for conserving the layer representation over consecutive predictive cycles. Conv: convolutional layer; BatchNorm: batch normalization layer; ReLU: rectifying linear unit; MaxPool: pooling layer; ConvTranspose: deconvolutional layer. The network received a 3-channel RGB image and a 1-channel depth image of the input object. The network output consisted of a 6-channel array, each channel representing a pixelwise map with the same dimensions as the input images and corresponding to one of the 6 grasp parameters: grasp centre coordinates (x, y); orientation (*θ*); grasp opening (h); gripper size (w); grasp quality (confidence) score. **(b)** Model performance during the inference phase, across 15 predictive coding timesteps, measured as the average mean squared error at each timestep, when presented with the test dataset that had been injected with varying levels of additive Gaussian noise. The right panel shows a representative sample object, corrupted with the different noise levels. **(c)** The average difference in test performance of the model between the initial (t=1) and final (t=15) timestep, for each noise level. Black bars denote standard errors of mean. **(d)** Left panel: an RGB image of a sample object from the test dataset. Right panel: the RGB image with Gaussian noise (*σ* = 0.25). **(e)** The grasp quality score maps of the sample object, at even timesteps.

**Table 1.** Neural network architecture of the short and long feedforward models. Layer-wise architecture of the feedforward models is described. All layers were implemented in PyTorch. The Conv2d (output channels, kernel size, stride) and ConvTranspose2d (output channels, kernel size, stride) refer to convolutional and deconvolutional layers, respectively. The MaxPool (kernel size, stride) layer implements feature pooling. BatchNorm2d: batch normalization; ReLU: rectifying linear unit; Tanh: hyperbolic tangent; Sigmoid: sigmoid function.

| Module |  | Layers | Short | Long |
| --- | --- | --- | --- | --- |
| Input | RGB | Conv2d (64, 3, 1) | • | • |
|  | Depth | Conv2d (64, 3, 1) | • | • |
| Feature compression |  | BatchNorm2d \| ReLU | • | • |
|  | Conv block 1 | Conv2d (64, 3, 1) \| BatchNorm2d \| ReLU | • | • |
|  |  | MaxPool (2, 2) | • | • |
|  | Conv block 2 | Conv2d (128, 3, 1) \| BatchNorm2d \| ReLU | • | • |
|  |  | Conv2d (128, 3, 1) \| BatchNorm2d \| ReLU | • | • |
|  |  | MaxPool (2, 2) | • | • |
|  | Conv block 3 | Conv2d (256, 3, 1) \| BatchNorm2d \| ReLU | • | • |
|  |  | Conv2d (256, 3, 1) \| BatchNorm2d \| ReLU | • | • |
|  |  | Conv2d (256, 3, 1) \| BatchNorm2d \| ReLU | • | • |
|  |  | MaxPool (2, 2) | • | • |
|  | Conv block 4 | Conv2d (256, 3, 1) \| BatchNorm2d \| ReLU | • |  |
|  |  | Conv2d (256, 3, 1) \| BatchNorm2d \| ReLU | • |  |
|  |  | Conv2d (512, 3, 1) \| BatchNorm2d \| ReLU |  | • |
|  |  | Conv2d (512, 3, 1) \| BatchNorm2d \| ReLU |  | • |
|  |  | Conv2d (512, 3, 1) \| BatchNorm2d \| ReLU |  | • |
|  |  | MaxPool (2, 2) |  | • |
|  | Conv block 5 | Conv2d (512, 3, 1) \| BatchNorm2d \| ReLU |  | • |
|  |  | Conv2d (512, 3, 1) \| BatchNorm2d \| ReLU |  | • |
|  |  | Conv2d (512, 3, 1) \| BatchNorm2d \| ReLU |  | • |
| Feature expansion |  | ConvTranspose2d (256, 3, 2) \| ReLU |  | • |
|  |  | Conv2d (256, 5, 1) \| ReLU |  | • |
|  |  | ConvTranspose2d (128, 3, 2) \| ReLU | • | • |
|  |  | Conv2d (128, 5, 1) \| ReLU | • | • |
|  |  | ConvTranspose2d (64, 3, 2) \| ReLU | • | • |
|  |  | Conv2d (128, 5, 1) \| ReLU | • | • |
|  |  | ConvTranspose2d (64, 3, 2) \| ReLU | • | • |
| Output | Grasp | ConvTranspose2d (5, 3, 1) \| Tanh | • | • |
|  | Confidence | ConvTranspose2d (1, 3, 1) \| Sigmoid | • | • |

#### Long feedforward model

The long backbone once again had a compression module followed by an expansion module similar to the short backbone, except, here the compression module comprised of a VGG16 model (Simonyan & Zisserman, 2014) with 5 convolution blocks where each convolution block contained two or three Conv layers with ReLU activation layers, followed by a MaxPool layer, and BN layers interspersed between each Conv and ReLU layer. Just like before the model architecture was modified with a two-arm input layer to receive 4-channel RGB-D input images. Also, the classification head of the canonical VGG16 architecture was replaced with the same regression head as the short backbone model to return the required parametric map output (see Table 1 for details).

### Predictive coding dynamics for feedback models

After training the feedforward backbone models, we augmented them with generative feedback connections adapting the PyTorch *Predify* library (Choksi et al., 2021) with some modifications that we made to the original code to introduce custom feedback connectivity. The library is designed to add predictive coding dynamics to existing deep neural networks. The predictive coding framework posits that the brain maintains an internal model of the world to actively predict sensory inputs (Rao & Ballard, 1999b). In this hierarchical model, higher areas generate predictions for lower areas, and discrepancies between predicted and actual inputs (prediction errors) are used to update and refine the higher-layer representations. This iterative process allows the network to minimize prediction errors and enhance perception of sensory information. For example, CNNs have been recently combined with feedback mechanisms for robust object perception (Choksi et al., 2021; Huang et al., 2020).

In the present study, we selected *N* encoding modules, 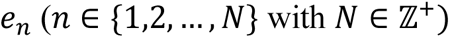 from the network backbone of the feedforward model and added *N* corresponding decoding modules, *d_n_*. An encoding module, *e_n_*, and the corresponding decoding module, *d_n_*, collectively constituted a *PCoder*. Each *e_n_* (except for e*_1_*) consisted of the MaxPool layer of a given convolution block and the two or three Conv/BN/ReLU layers of the subsequent convolution block. The output of *e_n_* was then passed through a feedback layer (deconvolution layer) *d_n_* that predicted the output of the last Conv/BN/ReLU layer of the previous convolution block (target). To this end, each *d_n_* consisted of a ConvTranspose2d layer that upscaled the input feature map to match the spatial dimensions of the target with feedback weights connecting module *n*+1 to module *n* being denoted by 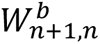. This way, when an input image initially activated all encoding modules through a feedforward pass, over subsequent successive recurrent iterations (timesteps *t*), both the decoding and encoding module representations were updated using the following equations:

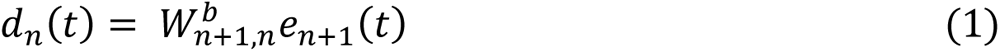

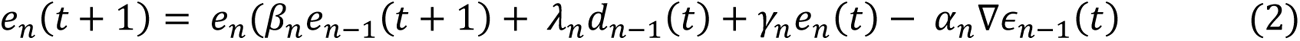

Here, 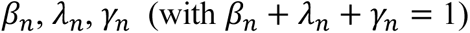 and *α_n_* served as balancing coefficients for the feedforward, feedback, recurrence and error-correction terms, respectively. The recurrence term, *γ_n_*, functioned as a memory buffer for retaining the encoding representation at the current timestep. The reconstruction error at module *n* − 1, denoted ∊*_n_*_–1_ (*t*), was defined as the mean squared error (MSE) between the feedforward representation e*_n_*_–1_ (*t*) and the predicted reconstruction *d_n_*_–1_ (*t*) at that timestep. The feedforward and feedback weights remained frozen across iterations.

#### Feedback models

Modifications were made to the original code from (Choksi et al., 2021) to introduce custom feedback connectivity to the forward models.

***Experiment 1.*** To test whether predictive coding aids not only object classification (Choksi et al., 2021; Huang et al., 2020) but also visuomotor processes, the network backbone of the short backbone model was augmented with 4 consecutive PCoders, similar to the feedback connectivity pattern attempted in (Choksi et al., 2021). The feedback connectivity for Experiment 1 is shown in Figure 1(a).

***Experiment 2.*** In order to identify the most effective source of feedback, feedback connections were implemented with a consistent feedback target layer and varying feedback source layers across model variants. Specifically, for each feedback type, 2 PCoders were designed. For Experiment 2, *e*_1_ was kept fixed as the first convolution block, across all feedback model variants. Notably, the feedback target layer for *e*_1_ was the input layer of the network; the input representations were configured to be static and were not updated by iterative recurrent mechanisms. The associated feedback loop was merely an appendage, which had no functional effect but was required in the model architecture for technical reasons. Additionally, the feedback source layer of *e*_2_ did not receive any recurrent feedback. Therefore, all model variants of feedback connectivity effectively contained a single functional loop of feedback, shown as solid red lines in Figure 2, for controlled comparisons between different feedback types.

**Figure 2.**
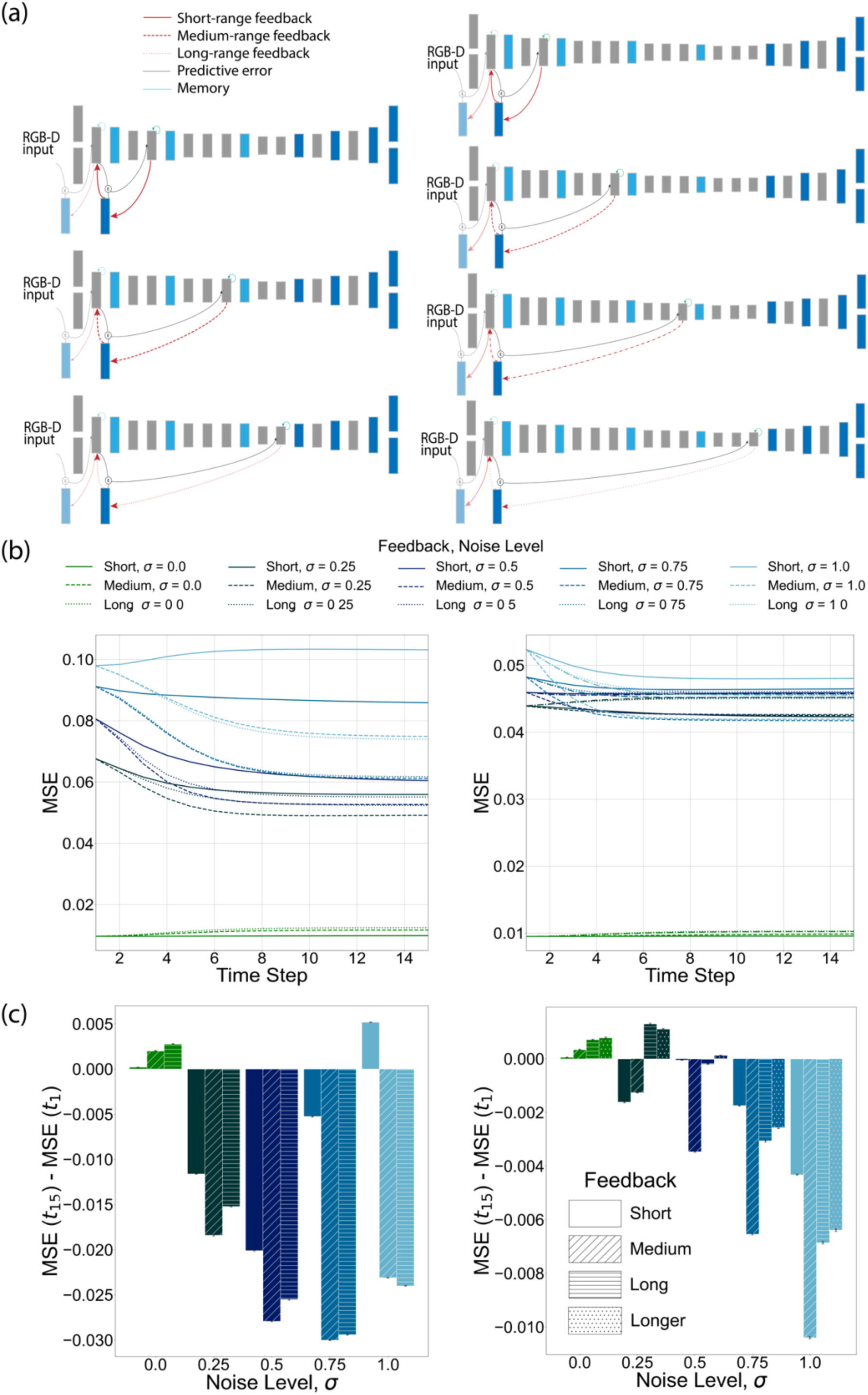
Layer-dependent effects of feedback with a uniform target. **(a)** Left panel: short feedforward model, augmented with *short*-range (first row; solid red arrow), *medium*-range (second row; dashed red arrow) and *long*-range (third row; dotted red arrow) feedback. Right panel: long feedforward model, augmented with *short*-range (first row; solid red arrow), *medium*-range (second row; dashed red arrow), *long*-range (third row; dash-dotted red arrow) and *longer*-range (fourth row; dotted red arrow) feedback. All effective feedback connections, pertaining to the second PCoder in each model variant, terminate at the same target layer: the final ReLU layer of convolution block 1. The first PCoder (translucent) does not represent a functional feedback connection. **(b)** Performance of feedback model variants across 15 predictive coding timesteps, on test dataset injected with varying levels of Gaussian noise (standard deviation, *σ* = {0, 0.25, 0.5, 0.75, 1.0}). Left panel: short feedforward backbone. Right panel: long feedforward backbone. **(c)** The average difference in test performance of the feedback model variants, between the initial (t=1) and final (t=15) timestep, for each noise level. Black bars denote standard errors of mean.

*Short backbone.* Three variants of feedback connectivity were compared: *short-*range, *medium-*range and *long-*range feedback, as shown in Figure 2(a). *e*_1_ comprised of the Conv/BN/ReLU layers of convolution block 1 for all model variants. *e*_2_ comprised of the following: the MaxPool layer of block 1 and the Conv/BN/ReLU layers of block 2 (*short-*range); the MaxPool layers of blocks 1 and 2, and the Conv/BN/ReLU layers of blocks 2 and 3 (*medium-* range); the MaxPool layers of blocks 1, 2 and 3, and the Conv/BN/ReLU layers of blocks 2,3 and 4 (*long-*range).

*Long backbone.* Four variants of feedback connectivity were compared: *short-*range, *medium-*range, *long-*range and *longer-*range feedback, illustrated in Figure 2(a). Similar to the short backbone, *e*_1_ comprised of the Conv/BN/ReLU layers of convolution block 1 for all model variants. *e*_2_ comprised of the following: the MaxPool layer of block 1 and the Conv/BN/ReLU layers of block 2 (*short-*range); the MaxPool layers of blocks 1 and 2, and the Conv/BN/ReLU layers of block 2 and 3 (*medium-*range); the MaxPool layers of blocks 1 ,2 and 3, and the Conv/BN/ReLU layers of block 2, 3 and 4 (*long-*range); the MaxPool layers of blocks 1, 2, 3 and 4, and the Conv/BN/ReLU layers of block 2, 3, 4 and 5 (*longer-*range).

***Experiment 3.*** To further isolate any effects of the feedback target layer, the feedback source layer was kept constant, while varying feedback target layers across model variants. Again, 2 PCoders were introduced into the feedforward network. Crucially, the feedback connection emanating from *e*_2_ was the only functional feedback loop. For all model variants, *e*_1_ spanned a single Conv layer that immediately precedes *e*_2_.

*Short backbone.* Three variants of feedback connectivity were compared: *short-*range, *medium-*range and *long-*range feedback. The feedback source layer of *e*_2_ was kept fixed as the final ReLU layer of convolution block 4 for all feedback variants. *e*_2_ comprised of the following: the MaxPool layer of block 3 and the Conv/BN/ReLU layers of block 4 (*short-*range); the MaxPool layers of blocks 2 and 3, and the Conv/BN/ReLU layers of blocks 3 and 4 (*medium-*range); the MaxPool layers of blocks 1,2 and 3, and the Conv/BN/ReLU layers of blocks 2, 3 and 4 (*long-*range). These model variants of feedback connectivity are shown in Figure 3(a).

**Figure 3.**
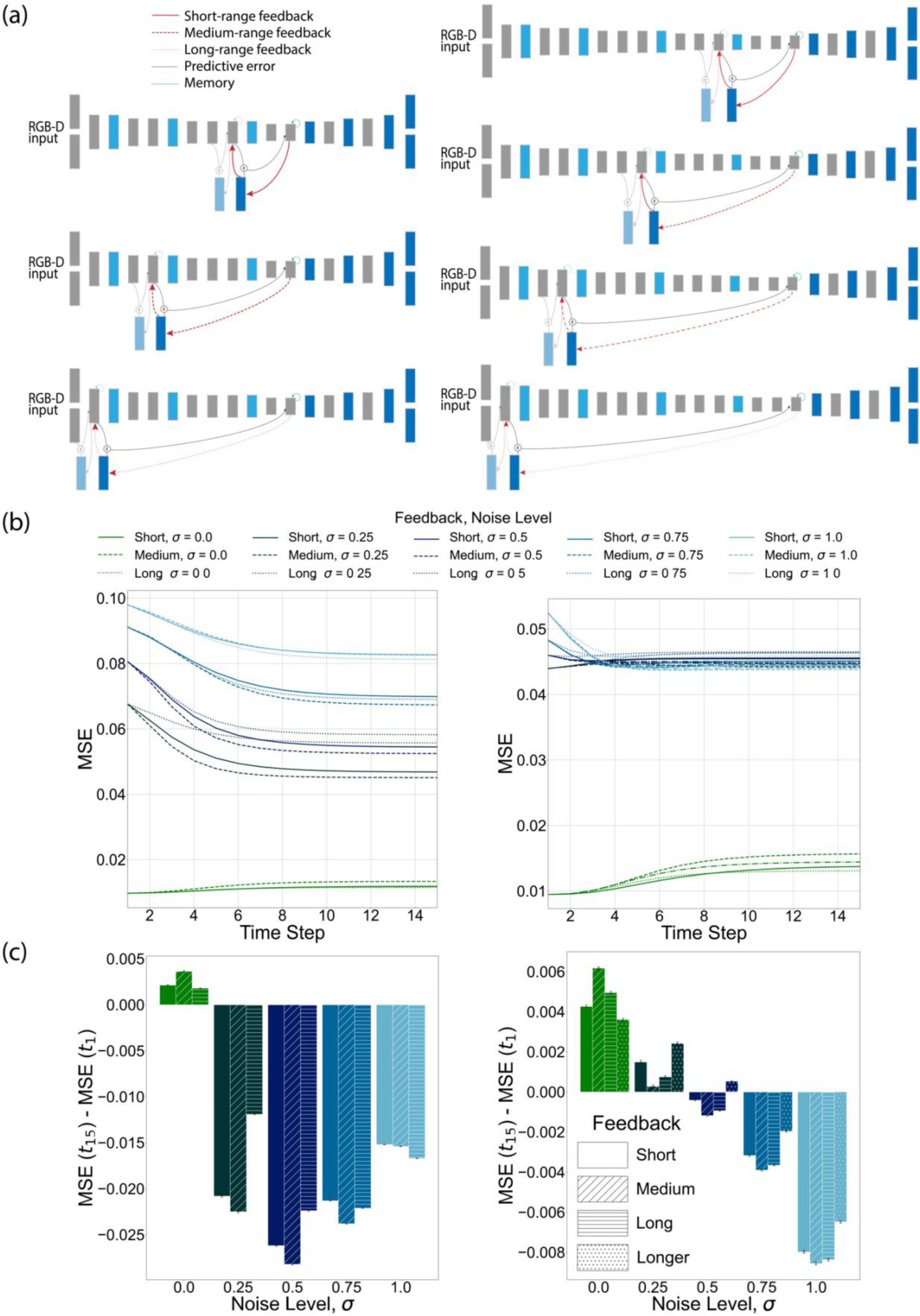
Layer-dependent effects of feedback with a uniform source. **(a)** Left panel: short feedforward model, augmented with *short*-range (first row; solid red arrow), *medium*-range (second row; dashed red arrow) and *long*-range (third row; dotted red arrow) feedback. Right panel: long feedforward model, augmented with *short*-range (first row; solid red arrow), *medium*-range (second row; dashed red arrow), *long*-range (third row; dash-dotted red arrow) and *longer*-range (fourth row; dotted red arrow) feedback. All effective feedback connections, pertaining to the second PCoder in each model variant, originate at the same source layer; short feedforward backbone: final ReLU layer of convolution block 4; long feedforward backbone: final ReLU layer of convolution block 5. The first PCoder (translucent) does not represent a functional feedback connection. **(b)** Performance of feedback model variants across 15 predictive coding timesteps, on test dataset injected with varying levels of Gaussian noise (standard deviation, *σ* = {0, 0.25, 0.5, 0.75, 1.0}). Left panel: short feedforward backbone. Right panel: long feedforward backbone. **(c)** The average difference in test performance of the feedback model variants, between the initial (t=1) and final (t=15) timestep, for each noise level. Black bars denote standard errors of mean.

*Long backbone.* Four variants of feedback connectivity were compared: *short-*range, *medium-*range, *long-*range and *longer-*range feedback. The feedback source layer of *e*_2_ was maintained as the final ReLU layer of convolution block 5 for all feedback variants. *e*_2_ comprised of the following: the MaxPool layer of block 4 and the Conv/BN/ReLU layers of block 5 (*short-*range); the MaxPool layers of blocks 3 and 4, and the Conv/BN/ReLU layers of blocks 4 and 5 (*medium-*range); the MaxPool layers of blocks 2, 3 and 4, and the Conv/BN/ReLU layers of blocks 3, 4 and 5 (*long-*range); the MaxPool layers of blocks 1, 2, 3 and 4, and the Conv/BN/ReLU layers of blocks 2, 3, 4 and 5 (*longer-*range). Model variants of feedback connectivity are shown in Figure 3(a).

***Experiment 4.*** The effect the abstraction level of the feedback source on model performance was also investigated. *Medium-*range feedback (i.e., the most effective feedback distance in Experiments 2 and 3) was implemented at three different levels of abstraction along the long backbone: *proximal*, *intermediate* and *distal* feedback loops. The naming convention is in relation to the distance from the input layer of the network. For each level of source abstraction, 2 PCoders were integrated into the network backbone. Like Experiment 2, feedback emerging from *e*_2_ was the only operant feedback connection, whereas the feedback loop across *e*_1_ traversed a single layer and had no predictive function.

For the three model variants, *e*_2_ comprised of the MaxPool layers of blocks 1 and 2, and the Conv/BN/ReLU layers of blocks 2 and 3 (*proximal*); the MaxPool layers of blocks 2 and 3, and the Conv/BN/ReLU layers of blocks 3 and 4 (only the first two Conv/BN/ReLU layers of block 4 were included in order to implement *medium-*range feedback distance) (*intermediate*); the MaxPool layers of blocks 3 and 4, and the Conv/BN/ReLU layers of blocks 4 and 5 (only the first two Conv/BN/ReLU layers of block 5 were included in order to maintain *medium-*range feedback distance) (*distal*). The model variants of medium-range feedback that were implemented are detailed in Figure 4(a).

**Figure 4.**
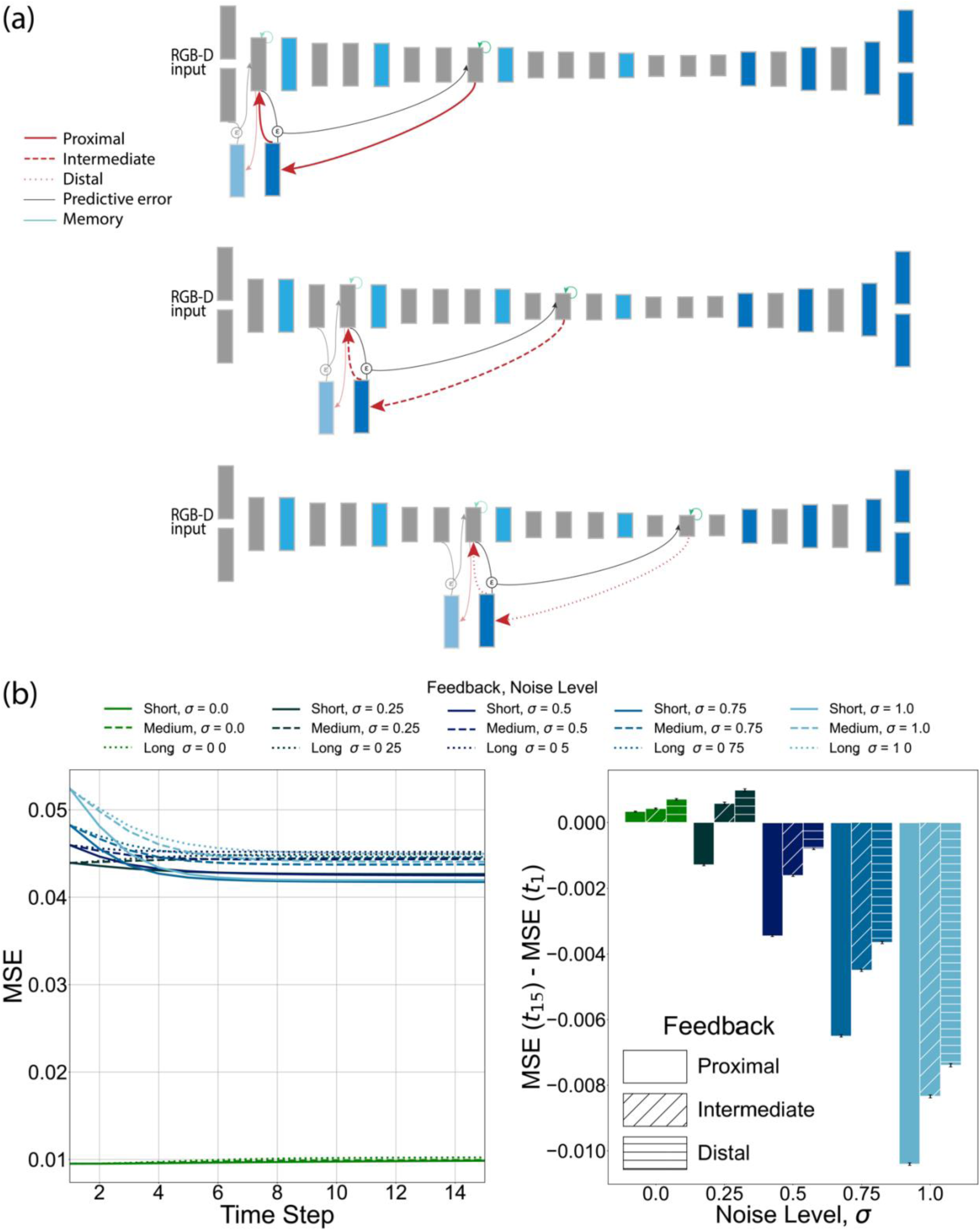
Effect of feedback source abstraction level. **(a)** Long feedforward model, augmented with *medium*-range feedback connections (pertaining to the second PCoder in each model variant), originating at layers *proximal* (first row; solid red arrow), *intermediate* (second row; dashed red arrow), or *distal* (third row; dash-dotted red arrow) to the input layer. The first PCoder (translucent) does not represent a functional feedback connection. **(b)** Left panel: performance of feedback model variants across 15 predictive coding timesteps, on test dataset injected with varying levels of Gaussian noise (standard deviation, *σ* = {0, 0.25, 0.5, 0.75, 1.0}). Right panel: the average difference in test performance of the feedback model variants, between the initial (t=1) and final (t=15) timestep, for each noise level. Black bars denote standard errors of mean.

### Model training and hyperparameters

Both, short and long backbone models were initialized with random weights (Xavier initialization) and trained on the task of grasp parameter estimation. Training involved error back-propagation and gradient descent to minimize the mean squared error (MSE) loss for the pixelwise regression output.

After hyperparameter optimization using a grid search strategy, both, the short and long forward models were trained using a Ranger optimizer, with a learning rate of 0.01, weight decay of 0, and a batch size of 75. The short and long backbone models were trained for 17 and 9 epochs, respectively.

The feedback models were created by adding recurrent feedback to the already trained backbone models. The activations of all encoding modules, *e_n_* were initiated with a feedforward pass. Then the forward weights were frozen, and the weights of the feedback deconvolution layers, *d_n_*, were trained using an unsupervised reconstruction objective. This seeks to minimize the reconstruction loss (prediction error), modelled as MSE between the outputs of *e_n−1_* and *d_n_*.

Hyperparameters for predictive coding dynamics, including layer-dependent balancing coefficients for the feedforward (*β_n_*), feedback (*λ_n_*), recurrence (*γ_n_*) and error-correction (*α_n_*) terms, were optimized using a grid search strategy. The respective hyperparameters were conserved across all model variants within each experiment and are presented in Table 2.

**Table 2.**
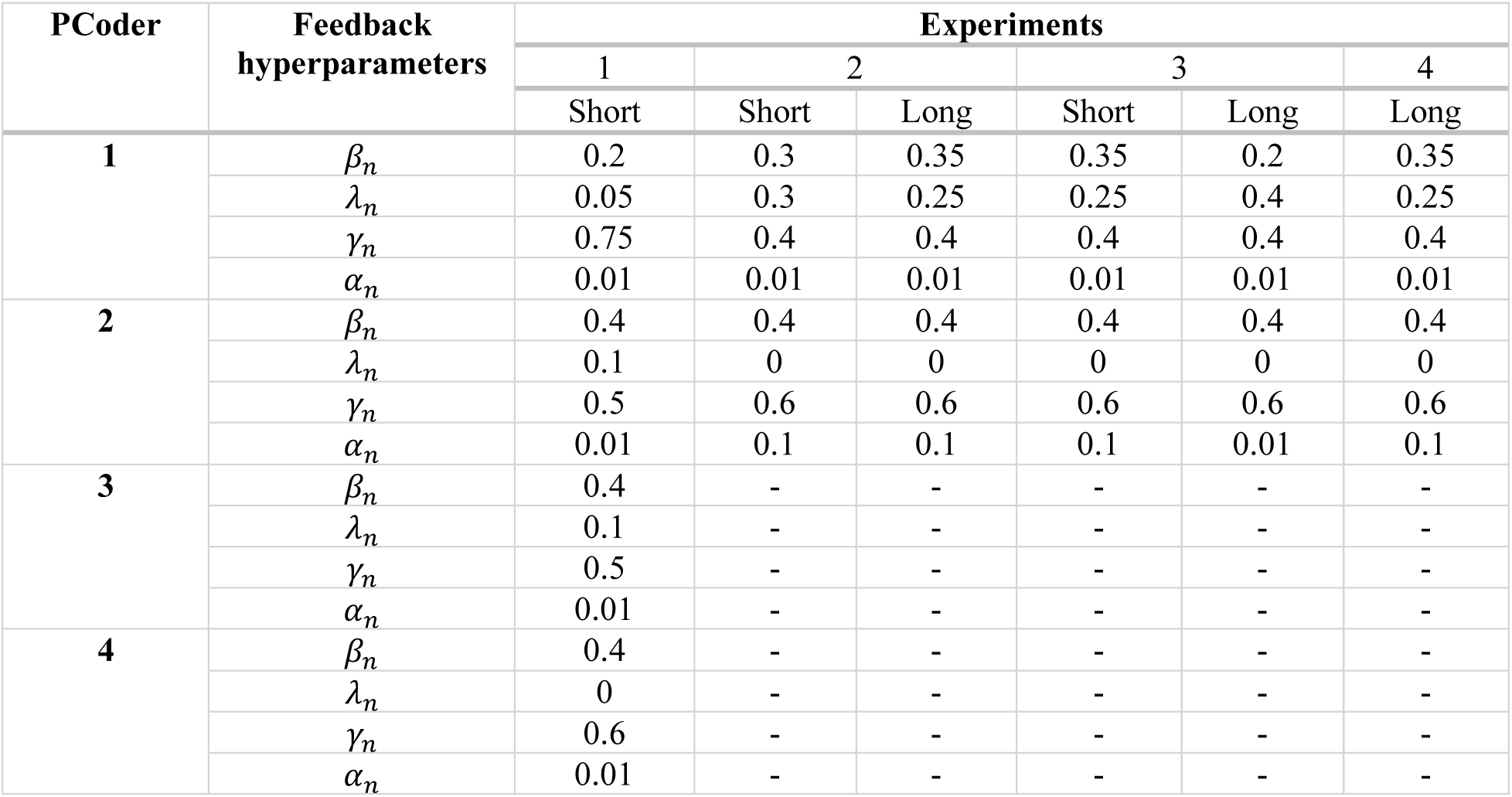
Predictive coding hyperparameters. The balancing coefficients for the feedforward (*β_n_*), feedback (*λ_n_*), recurrence (*γ_n_*) and error-correction (*α_n_*) terms that were selected after hyperparameter optimization, are stated for the short and/or long feedforward model(s), for all experiments.

| PCoder | Feedback hyperparameters | Experiments |  |  |  |  |  |
| --- | --- | --- | --- | --- | --- | --- | --- |
|  |  | 1 | 2 |  | 3 |  | 4 |
|  |  | Short | Short | Long | Short | Long | Long |
| **1** | $\beta_n$ | 0.2 | 0.3 | 0.35 | 0.35 | 0.2 | 0.35 |
| | $\lambda_n$ | 0.05 | 0.3 | 0.25 | 0.25 | 0.4 | 0.25 |
| | $\gamma_n$ | 0.75 | 0.4 | 0.4 | 0.4 | 0.4 | 0.4 |
| | $\alpha_n$ | 0.01 | 0.01 | 0.01 | 0.01 | 0.01 | 0.01 |
| **2** | $\beta_n$ | 0.4 | 0.4 | 0.4 | 0.4 | 0.4 | 0.4 |
| | $\lambda_n$ | 0.1 | 0 | 0 | 0 | 0 | 0 |
| | $\gamma_n$ | 0.5 | 0.6 | 0.6 | 0.6 | 0.6 | 0.6 |
| | $\alpha_n$ | 0.01 | 0.1 | 0.1 | 0.1 | 0.01 | 0.1 |
| **3** | $\beta_n$ | 0.4 | - | - | - | - | - |
| | $\lambda_n$ | 0.1 | - | - | - | - | - |
| | $\gamma_n$ | 0.5 | - | - | - | - | - |
| | $\alpha_n$ | 0.01 | - | - | - | - | - |
| **4** | $\beta_n$ | 0.4 | - | - | - | - | - |
| | $\lambda_n$ | 0 | - | - | - | - | - |
| | $\gamma_n$ | 0.6 | - | - | - | - | - |
| | $\alpha_n$ | 0.01 | - | - | - | - | - |

Model training was performed on the NVIDIA T4 and the NVIDIA QUADRO RTX 8000 GPUs.

### Dataset

Both, the backbone and feedback models, were trained on the grasp detection task using the large-scale Jacquard dataset for robotic grasp detection (Depierre et al., 2018). The Jacquard dataset is composed of 11,619 distinct objects with a total of ∼5,000,000 possible grasp annotations. The grasp annotations corresponding to each image constituted a list of grasp candidate labels. Each grasp candidate was described by parameters including center pixel coordinates (x, y), orientation (theta), and dimensions (pixel width, pixel height) of the grasp. For each object instance (a distinct camera viewpoint of an object), a rendered RGB image, a segmentation mask, two depth images and the grasps annotations are available. The dataset was split into three parts: 66% for model training, 17% for model validation, and 17% for model testing.

#### Preprocessing

The data preprocessing pipeline transformed lists of grasp candidates into pixel-wise grasp maps. A grasp map array, of spatial dimensions corresponding to the image size, was initialized with six channels to represent the five grasp parameters and a count of overlapping grasp bounding boxes for each pixel. The preprocessing operation converted grasp annotations to bounding boxes, identified the bounding box boundaries, and iterated over each pixel within these boundaries to determine the grasp candidate closest to the pixel based on Euclidean distance. Each pixel was then labeled with the attributes of the nearest grasp and the count of valid bounding boxes covering it. The final channel, representing the count of bounding boxes per pixel, was normalized by dividing by the maximum count across all pixels. This preprocessed data format was used for training the backbone and feedback neural network models on the task of predicting grasp locations in an input image.

Feature normalization was applied to all images (RGB and depth) and grasp labels, to promote training stability and model convergence. Image pixel intensities and label values were normalized to a range between 0 and 1.

#### Additive noise

During model inference, all test images were injected with varying levels of additive Gaussian noise of standard deviation, *σ* ∈ {0, 0.25, 0.5, 0.75, 1}.

### Statistics

A repeated measures analysis of variance (ANOVA) (Girden, 1992) was conducted to assess the effects of feedback type and noise level on the grasp detection accuracy of the model, using the Jamovi statistical software (The jamovi project, 2024). This analysis accounted for within-object effects and included tests for sphericity. Mauchly’s test of sphericity (Mauchly, 1940) was performed to check the assumption of sphericity for feedback type, noise level, and their interaction. Where the assumption was violated, Greenhouse-Geisser corrections (Greenhouse & Geisser, 1959) were applied to adjust the degrees of freedom. Post hoc comparisons were conducted using Tukey’s HSD test (Tukey, 1949) to evaluate pairwise differences between the levels of feedback type.

A One-Way ANOVA (David C. Howell, 2002) was conducted to compare the difference in model performance between the first and final timestep during the test, across noise different noise levels (The jamovi project, 2024).

### Code Availability

All code used for this study is available at https://github.com/khanrom/Feedback_Grasping.

## Results

We investigated the contribution of feedback to vision for action, such as grasp planning, using a neural network model that was optimized to detect grasp points on objects.

### Feedback improves robustness of sensorimotor control to noise

We evaluated the effect of predictive feedback on the performance of the trained model, when presented with adversarial images that were injected with varying levels of Gaussian noise (*σ* = {0, 0.25, 0.5, 0.75, 1.0}) (Experiment 1 in Methods). In effect, the performance of a feedback model at the first timestep (*t* = 1) of the iterative predictive coding dynamics is identical to the performance of the corresponding feedforward backbone. This backbone performance was compared to the performance of the feedback model after several recurrent timesteps (*t* = 15) after which performance no longer changed. We observed an improvement in the grasp detection accuracy of the model after predictive coding iterations, across all levels of Gaussian noise (Figure 1B; n.b., there was a small decline in performance for zero noise). The average difference in model performance between the first (*t* = 0) and the final (*t* = 15) timestep was significant across noise levels, *F*(4, 22267) = 220036.00, *p* < .001 (Figure 1C). Figure 1E depicts progressive noise removal in the model output (represented by the grasp quality score map) across successive predictive coding timesteps, for a sample object from the test dataset (Figure 1D).

### Performance-related effects of feedback are layer-dependent

Next, we tested whether the performance-enhancing effects of feedback were contingent on layer-specific feedback connectivity. To probe this question, we contrasted different patterns of feedback connectivity.

#### Constant feedback target

First, we set up model variants augmented with a constant feedback target (final ReLU layer of convolution block 1) that received *short*-, *medium*-, or *long*-range feedback, for the short backbone and an additional *longer*-range feedback variant for the long backbone, as detailed in the Methods (see Experiment 2). The average difference in model performance between the first (t=0) and the final (t=15) for all feedback model variants was compared.

For the short backbone, *medium*-range feedback confers the largest advantage on model performance on average under Gaussian noise (*σ* = {0, 0.25, 0.5, 0.75}), as seen in Figure 2(b,c). A repeated measures ANOVA after Greenhouse-Geisser correction yielded a significant main effect of noise-dependent benefit of feedback on model performance (see Table 3) reflecting that performance declined with increasing noise, as expected. More importantly, there was a significant main effect of feedback distance (see Table 4). Post hoc comparisons using Tukey’s HSD test revealed significant differences in model performance between all pairs of feedback types with medium-range feedback being the most effective on average (see Table 6). Additionally, there was a significant interaction between feedback distance and noise level (see Table 5), indicating that most effective feedback distance depended on the noise-dependent benefit (*medium*-range feedback for intermediate levels of noise, *short*-range for zero noise, *long*-range for high levels of noise).

**Table 3.** Within Subjects Effects - Noise-dependent benefit.

| **Experiment** |  |  | **df** | **F** | **p** | **$\eta^2$** |
| --- | --- | --- | --- | --- | --- | --- |
| **2** | Short backbone | Noise-dependent benefit | 1.68 | 109649 | < .001 | 0.454 |
|  |  | Residual | 15772.36 |  |  |  |
|  | Long backbone | Noise-dependent benefit | 1.64 | 43410 | < .001 | 0.426 |
|  |  | Residual | 15398.52 |  |  |  |
| **3** | Short backbone | Noise-dependent benefit | 1.56 |  |  |  |
|  |  | Residual | 14671.10 | 59300 | < .001 | 0.652 |
|  | Long backbone | Noise-dependent benefit | 2.71 | 24519 | < .001 | 0.376 |
|  |  | Residual | 25388.71 |  |  |  |
| **4** | Long backbone | Noise-dependent benefit | 2.13 | 48395 | < .001 | 0.520 |
|  |  | Residual | 19937.27 |  |  |  |

**Table 4.** Within Subjects Effects - Feedback distance (abstraction level)

| Experiment | | | df | F | p | $\eta^2$ |
| --- | --- | --- | --- | --- | --- | --- |
| 2 | Short backbone | Feedback distance | 1.18 | 87764 | < .001 | 0.194 |
|  |  | Residual | 11072.25 |  |  |  |
|  | Long backbone | Feedback distance | 1.64 | 10542 | < .001 | 0.080 |
|  |  | Residual | 15429.00 |  |  |  |
| 3 | Short backbone | Feedback distance | 1.10 | 5365 | < .001 | 0.010 |
|  |  | Residual | 10297.82 |  |  |  |
|  | Long backbone | Feedback distance | 2.61 | 2908 | < .001 | 0.004 |
|  |  | Residual | 24479.40 |  |  |  |
| 4 | Long backbone | Feedback abstraction level | 1.35 | 13523 | < .001 | 0.039 |
|  |  | Residual | 12629.91 |  |  |  |

**Table 5.**
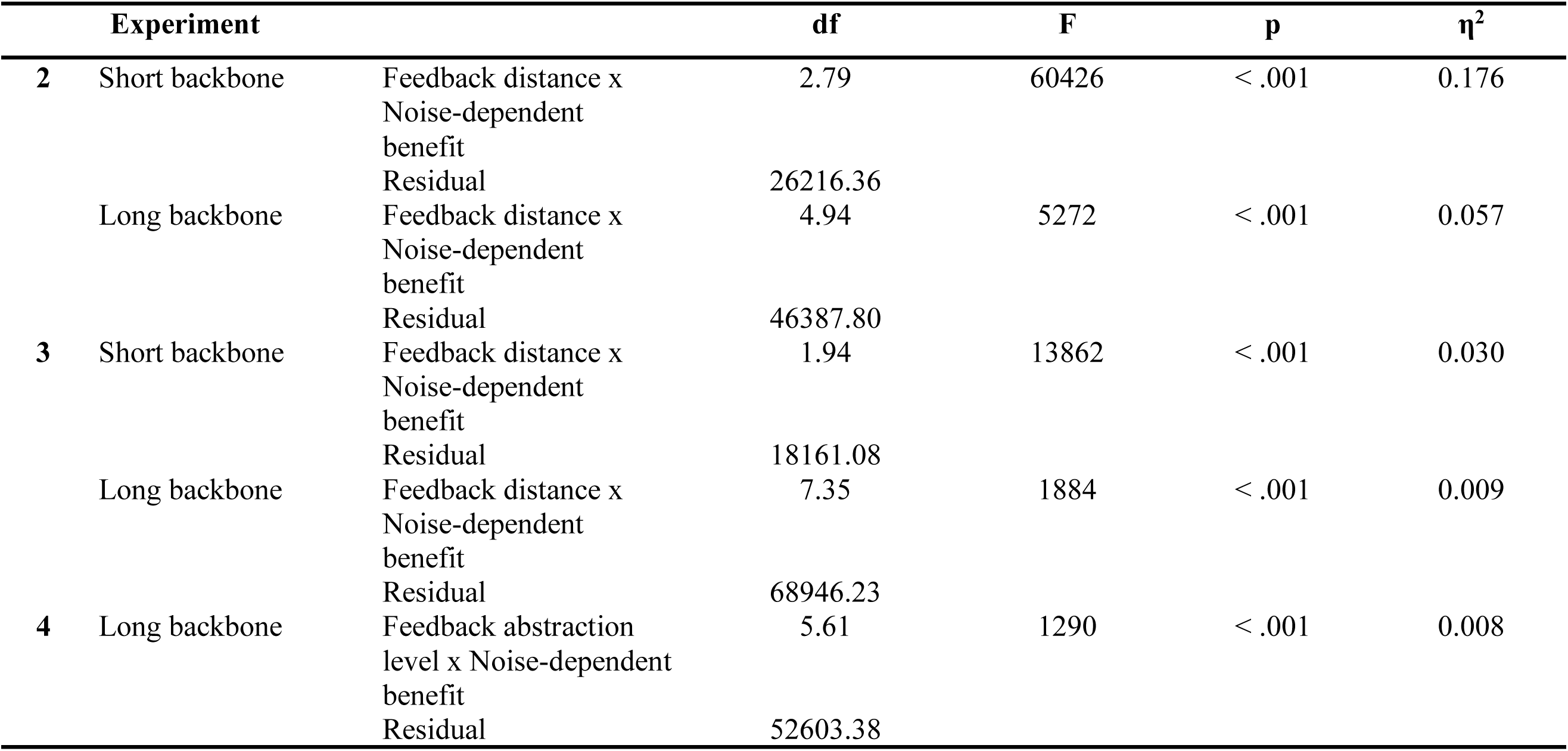
Within Subjects Effects - Feedback distance (abstraction level) x Noise-dependent benefit.

**Table 6.**
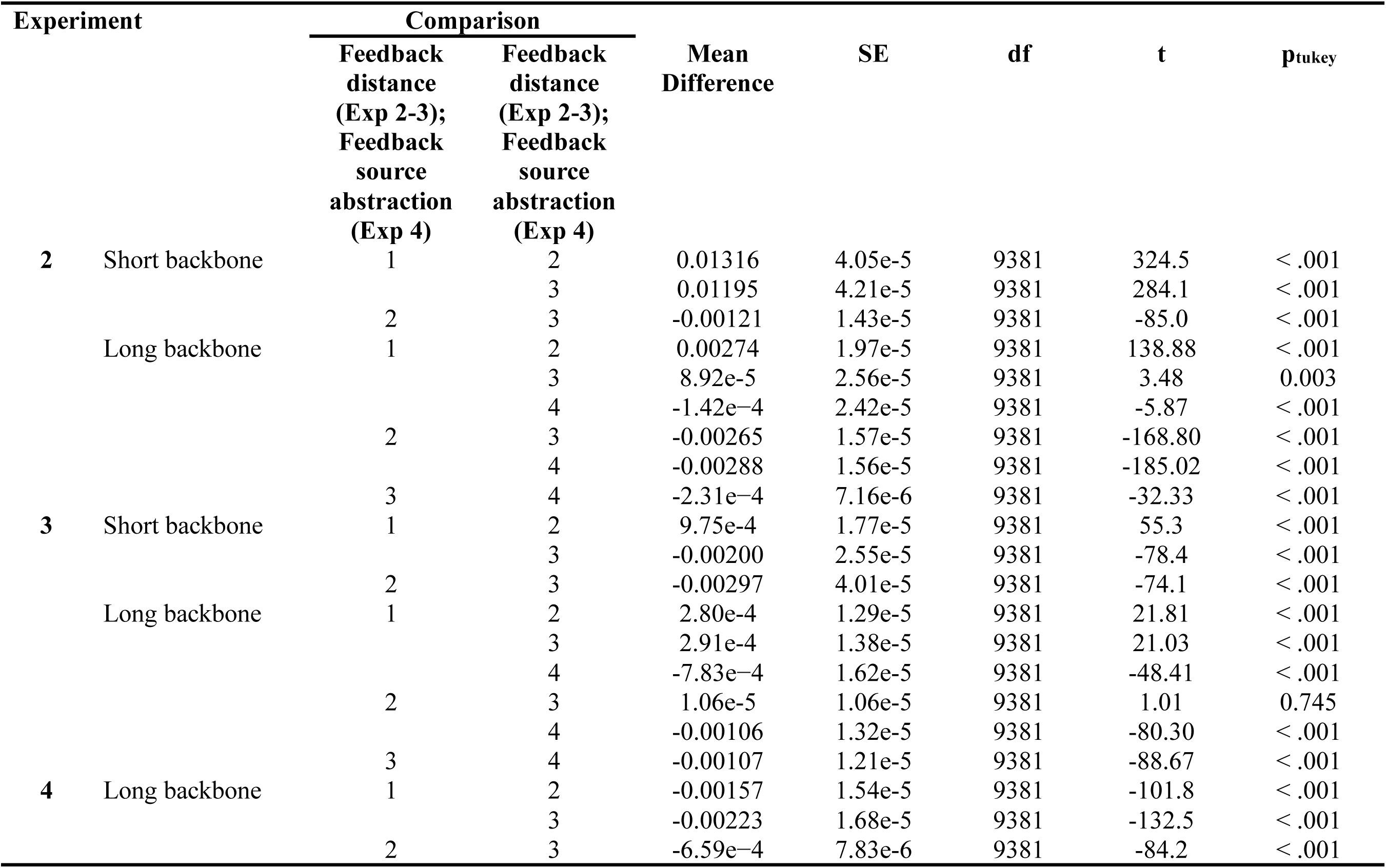
Post Hoc Comparisons - Feedback distance (abstraction level)

Similarly, in the case of the long backbone, *medium*-range feedback showed on average the greatest robustness to noise (*σ* = {0.5, 0.75, 1.0}), as shown in Figure 2(b,c). A Greenhouse-Geisser corrected repeated measures ANOVA showed significant main effects of noise level (see Table 3), and feedback distance (see Table 4), where Tukey’s HSD test confirmed that *medium*-range feedback was most effective establishing significant differences between various pairs of feedback types (see Table 6). Once again, the interaction between feedback distance and noise-dependent feedback benefit was also significant (see Table 3), indicating that *medium*-range feedback was most effective for intermediate and high levels of noise whereas *short*-range feedback was more effective for zero and low levels of noise.

#### Constant feedback source

Secondly, we compared model performance under adversarial noise, when the feedback connections originated from the same source layer across all model variants (for details of the model architecture, see Experiment 3 in Methods). Model variants were evaluated after timestep *t* = 15. Once again for the short backbone we found *medium*-range feedback was most ameliorative, when presented with stimuli injected especially with intermediate levels of Gaussian noise (*σ* = {0.25, 0.5, 0.75}), as shown in Figure 3(b,c). A repeated measures ANOVA with Greenhouse-Geisser correction demonstrated a significant main effect of noise-dependent benefit of feedback (see Table 3). More importantly, we observed a significant main effect of feedback distance on model performance (see Table 4), with *medium*-range feedback yielding the greatest benefit on average. In detail, Tukey’s HSD test revealed significant differences between various pairs of feedback types (see Table 6). Finally, there was a significant interaction between feedback distance and noise-dependent benefit (see Table 5), , indicating that *medium*-range feedback was optimal for intermediate levels of noise whereas *long*-range feedback was more ameliorative for high and zero levels of noise.

Similar effects of feedback connectivity on model performance were observed in the case of the long backbone, Figure 3(b,c). The network augmented with *medium*-range feedback was most resilient to all levels of additive noise tested, except for zero noise (*σ* = {0.25, 0.5, 0.75, 1.0}). A repeated measures ANOVA with Greenhouse-Geisser correction showed significant main effects of noise-dependent feedback benefit (see Table 3), and feedback distance (see Table 4). Post hoc comparisons using Tukey’s HSD test revealed significant differences between the various pairs of feedback types (see Table 6). These results are consistent with *medium*-range feedback being most beneficial across noise levels on average. Furthermore, a significant interaction between feedback distance and noise-dependent benefit of feedback (see Table 6) reflected that *medium*-range feedback was optimal for most noise levels except zero noise.

### Proximal neural feedback loops improve robustness to noise

Having established the precedence of medium-range neural feedback loops in two distinct neural network architectures, we investigated if the performance-enhancing effects of such feedback were sensitive to the level of abstraction of the feedback source, i.e., feedback originating from early vs. late areas in the dorsal visual stream. Using the long backbone architecture, we set up three different model variants, augmented with *medium*-range feedback connections, originating at *proximal*, *intermediate* or *distal* layers relative to the input (see Experiment 4 in Methods), which were then presented with input images corrupted with Gaussian noise. Feedback most *proximal* to the input layer, had the greatest performance-enhancing effect, as seen in Figure 4(b). A repeated measures ANOVA with Greenhouse-Geisser correction examined the effects of feedback source abstraction level, as well as noise level on the change in performance of the model between timestep *t* = 0 and timestep *t* = 15. The analysis identified significant main effects of noise-dependent benefit of feedback (see Table 3), as well as feedback abstraction level (Table 4). The latter effect reflected proximal feedback to be optimal as shown with post hoc comparisons using Tukey’s HSD test indicating significant differences between various pairs of feedback abstraction levels (see Table 6). Moreover, the interaction between abstraction and noise level was significant (see Table 5) due to the influence of abstraction level being less prominent for zero noise than for other noise levels.

## Discussion

In the present study, we investigated the contribution of neural feedback to sensorimotor functions, in particular visual processing during grasp planning, by utilizing convolutional neural network models that had been augmented with predictive feedback and that were trained to compute grasp positions for real-world objects. After establishing an ameliorative effect of symmetric feedback on grasp detection from noisy images, we characterized the performance effects of asymmetric feedback, similar to that observed in the cortex. We found that the performance-enhancing effect of predictive coding under adverse conditions was optimal, across a range of noise levels, for *medium*-range asymmetric feedback. Moreover, this effect was most prominent when *medium*-range feedback originated at a level of representational abstraction that was proximal to the input layer, in contrast to more distal layers.

Prior computational work has demonstrated that symmetric feedback, originating from the immediate next higher area in the cortical hierarchy, helps suppress irrelevant noise to enhance performance in an object classification task (Choksi et al., 2021; Huang et al., 2020). Importantly, our work extends the role of predictive feedback in noise-cleanup to the more general function of sensorimotor processing. This involves progressively denoising the feature representations during the test phase, essentially by projecting the representations onto the learned manifolds in the representational space of the trained model. Consistent with this idea, iterative predictive coding dynamics worsen the model’s performance when presented with clean images, possibly because the model, after finding a suitable solution with the first forward pass, converged onto a less optimal average solution within the latent representational space.

However, accumulating neurophysiological evidence has suggested that cortical feedback pathways are not strictly reciprocal (Salin & Bullier, 1995) and may traverse multiple areas at different latencies (Bullier, 2001; Markov et al., 2014). By systematically comparing different feedback path lengths and levels of representational abstraction, we illustrated that too short a feedback loop may lack the broader contextual information needed to mitigate noise effectively, while longer loops may carry overly abstract predictions. We propose that *medium*-range feedback marks a unique sweet spot, putatively incorporating an optimal neural transmission delay and an informative level of abstraction of the feedback source. Moreover, our results suggest that *medium*-range feedback benefits noise clean-up the most when it originates from layers that are proximal to the input, suggesting that relatively early representations are most effective in removing noise.

Of note is the observed U-shaped effect of noise-dependent performance benefit of predictive feedback for the short backbone experiments (Fig 2C, left; Fig3C, left) vs. a linear decrease observed for the long backbone models (Fig 2C, right; Fig3C, right; Fig 4B, left), across the five discrete levels of additive Gaussian noise *σ* = {0, 0.25, 0.5, 0.75, 1.0} that were tested. We speculate that the linear trend seen for the long backbone is the arm of a shifted, wider U-shape reflecting robustness to noise perturbation and recovery for the higher-dimensional (in terms of parametric space) long backbone models. The hypothesized U-shaped may emerge if the long backbone model variants are evaluated on higher levels of additive Gaussian noise e.g. *σ* = {1.25, 1.5, 1.75, … }.

Together our findings suggest a previously uncharted role of intermediate cortical areas along the dorsal stream, e.g., roughly similar to area V6A in the primate brain, as a source of predictive feedback. V6A is primarily involved in encoding action-guided vision during grasping behaviors (Galletti & Fattori, 2018). It plays a key role in guiding prehension movements (Galletti et al., 2003). Importantly, V6A integrates both visual inputs and somatosensory feedback from the upper limbs (Gamberini et al., 2011). Our results indicate that V6a might be at a strategically optimal position to exert dynamic top-down influence onto early visual areas during visuomotor processing, this way constituting an important hypothesis for future research. although the extent to which the present neural networks exhibit V6a-like properties requires further investigation.

Also intriguing is the observation that feedback arising from proximal layers of the network is more beneficial than feedback coming from distal layers. Interestingly, the opposite pattern of effectiveness was reported for simulated feedback emulating attentional effects in a VGG16 network (Lindsay & Miller, 2018). Lindsay and Miller (2018) found that attentional effects were most noticeable when injected into more distal layers. This difference in proximal vs. distal efficacy of feedback is not explained by structural differences in the feedforward networks given that the current study used the same long backbone employed previously (Lindsay & Miller, 2018). The difference in efficacy could be due to task differences. However, Lindsay and Miller (2018) observed the same distal benefits a range of tasks. Instead, the proximal vs. distal differences might, and probably do, come from the disparity in feedback.

Whereas in the present study we implemented predictive feedback signals, Lindsay and Miller (2018) simulated attentional feedback according to the feature similarity gain model (Martinez-Trujillo & Treue, 2004; Treue & Trujillo, 1999) where neuronal activity changes as a function of similarity of a neuron’s tuning to an attended feature. Whether this particular kind of attentional modulation or more generally the function of attention as a form of task-relevant resource allocation in contrast to task-independent predictive coding (Kok, Rahnev, et al., 2012) can explain the differences between proximal vs. distal effectiveness should be explored in the future.

In conclusion, our simulations show that introducing biologically realistic asymmetric predictive feedback improves model robustness to noisy visual stimuli in a neural network model optimized for sensorimotor transformations. Our findings highlight the possible functional significance of mid-tier dorsal stream areas (e.g., V6A) that lie between low-level (V1) and high-level (parietal) representations, suggesting these areas might serve as ideal “predictors” for early visuomotor processing.

## Acknowledgement

This research was supported in part by a grant from the Natural Sciences and Engineering Research Council of Canada (NSERC).

